# Optimization of Alginante-poly-L-lysine Microencapsules Strength by Box-Behnken Model for CHO cells Culture in Stirred Tank Bioreactor

**DOI:** 10.1101/165464

**Authors:** Yu Wang, Na Li, Dongsheng Sun, Shengnan He, Lisha Mou

## Abstract

**Summary statement:** Box-Behnken model is an efficient method to optimized process conditions of microencapsules strength for long-term cell culture, promotes CHO cells viability and protein production in a stirred tank bioreactor.

**ABSTRACT:** Cell microencapsulation technology has been proved to be a valuable technology in the fields of large-scale cell culture. It is important to accurately construct the microcapsule membranes with desired properties including a certain thickness suitable for cell growth, and maximum strength for the stability of microencapsules. As single factor experiments are time-consuming to obtain the desired membrane preparation conditions, Box-Behnken model was used to investigate the interactions among reaction conditions, predict the optimized reaction conditions for given purpose, that was membrane with maximum strength and desired thickness for microencapsulated cell culture. Significant values of R^2^ in this study indicated the model theoretical values are very close to the measured values. Based on the desired membrane thickness, the process of maximum strength was optimized, and the prediction agreement of measured value and model theoretical value was 91.11%. The optimized microencapsules with maximum strength and 15 μm membrane thickness promote CHO cells viability and protein production in stirred tank bioreactor. The result shows that Box-Behnken model is an efficient method to optimized process conditions of microencapsules strength for long-term cell culture.

## Introduction

Cell microencapsulation technology has been proved to be a valuable technology in the fields of large-scale cell culture (Lim and SUN, 1980), enzyme immobilization (Cheng and Sefton, 2009) and artificial cells/organs (Luo et al., 2007). Alginante-poly-L-lysine (Alg-PLL) is one of the most well-studied microcapsules because of its good biocompatibility and good characteristics in cell culture (Liu et al., 2017; Domżalska et al., 2016). Alg-PLL microencapsulation provides the cells with polymeric semi-permeable membrane and a unique environment for high cell density and high productivity of proteins in mammalian cell culture (Zhang et al., 2008). Microencapsulation membrane allows bidirectional diffusion of nutrients, oxygen, and metabolic wastes, and prevents mammalian cells from destroying of stir (Fernandesa et al., 2007).

Membrane performance plays a key role on microencapsule’s strength and permeability (Zheng et al., 2016), which is highly related to membrane thickness. The microcapsules strength determined the stability of microencapsules during the preparation process and long-term culture, while membrane thickness effects the viability of microencapsulated cells. High membrane strength is required to resist the stresses and keep the stability of microencapsules during the preparation process and long-term cell culture. Membrane thickness is highly related to membrane strength, thicker membrane protects cells better with higher strength (Fernandesa et al., 2007; Ma et al.,1994). However, thicker membrane often results in lower mass transfer coefficient and further poorer permeability for nutrients, which limits the cell growth (Breguet et al.,2007; Goosen et al.,1989; Ooijkaas et al.,1999; Pajić-Lijaković et al., 2007).

As carriers for long-term cell culture, membrane of microencapsules is required with high strength while the long-term microencapsulated cells culture. The stress on the membrane of microencapsules comes from hydrogels swelling, cell cluster enlarging inside (Fernandesa et al., 2007) and shear stress outside. Previous research has demonstrated that microencapsulated recombinant CHO increased the DSPA (Desmodusrotundus salivary plasminogen activator) production in stirred tank bioreactor (Wang et al., 2015). However, it was found amount of microencapsules were broken by long-time string. Membrane strength is also crucial for microencapsulated islets implantation, which has been applied in clinical for diabetes treatment. Microencapsules with low strength would be destroyed by the blood fluid in vivo and pressure between organs. In our research, 15 μm was determined as appropriate membrane for a long-time cell culture (Ma et al., 2013). Therefore, it is necessary to optimize the membrane strength with specific thickness.

Box-Behnken design is a mathematical and statistical technique for building empirical models for optimizing production conditions, such as culture conditions (King et al., 1987; Saelao et al., 2011; Adinarayana and Ellaiah, 2002; Liu et al., 2003; Singh et al., 2009), enzymes production (Ismael et al., 1998; Park et al., 2002), and biomass production. Reduces the number of assays to optimize the process and gathers results more precise than those obtained by unilabiate strategies (Dutta et al., 2017), overcomes the shortcoming of the rational orthogonal method. Box-Behnken design could build models of several response values at the same time and optimize one of them with other response values are limited.

In this study, we optimized the preparation conditions of microcapsule membranes with Box-Behnken design. Two models built with Box-Behnken which was used to optimize the preparation conditions of microcapsules according to the responses to membrane thickness and swelling degree. Each of response values could be optimized as the other one was limited. Both models can provide more reliable predictive data for membrane preparation, and each of them so that the certain microcapsules can be obtained with desired membrane thickness and high membrane strength. According to the model, microencapsulated CHO cells with membrane thickness of 15 μm and maximum strength were prepared and cultured in stirred tank bioreactor.

## Results

### Results of Box-Behnken methods for screened

Calculated Sw according formula 1, denoted as Y1. Membrane thickness was recorded and denoted as Y2. The results were shown in Tab 1.

**Tab 1.** Test results of Box-Behnken Design.

| Reaction conditions | Y<sub>1</sub> (%) | Y<sub>2</sub>(μm) |
| --- | --- | --- |
| 1 | 80.84 | 5 |
| 2 | 83.33 | 10 |
| 3 | 50.77 | 10 |
| 4 | 40.71 | 15 |
| 5 | 23.27 | 10 |
| 6 | 96.88 | 5 |
| 7 | 13.40 | 15 |
| 8 | 13.03 | 15 |
| 9 | 12.85 | 15 |
| 10 | 96.62 | 5 |
| 11 | 39.24 | 15 |
| 12 | -13.50 | 20 |
| 13 | 10.71 | 15 |
| 14 | 8.93 | 15 |
| 15 | -23.25 | 20 |

2 Y<sub>1</sub>, swelling degree; Y<sub>2</sub>, membrane thickness.

### Factors influencing membrane strength and thickness

The values of S_w_ calculated above were fitted into model by Design expert. The polynomial models for response Y1 (Sw) and Y2 (membrane thickness) could be represented by following equations:

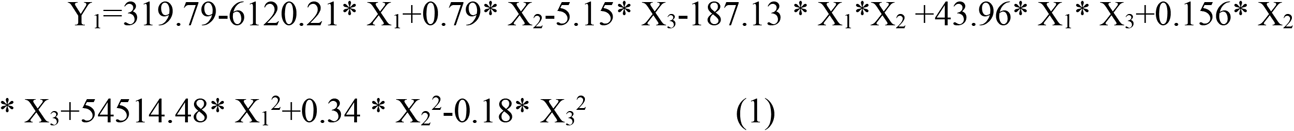

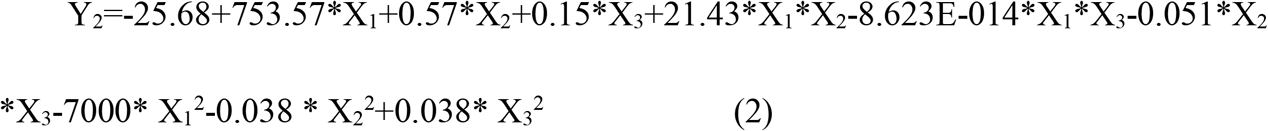

Fig. 1 showed the actual values for the response and predicted values determined by the model equations. The ANOVA for quadratic regress models was performed and was summarized in Tab 2. The results of model summary showed the R^2^ value was 0.9476 and 0.9454 separately for Y_1_ and Y_2_.

**Fig. 1.**
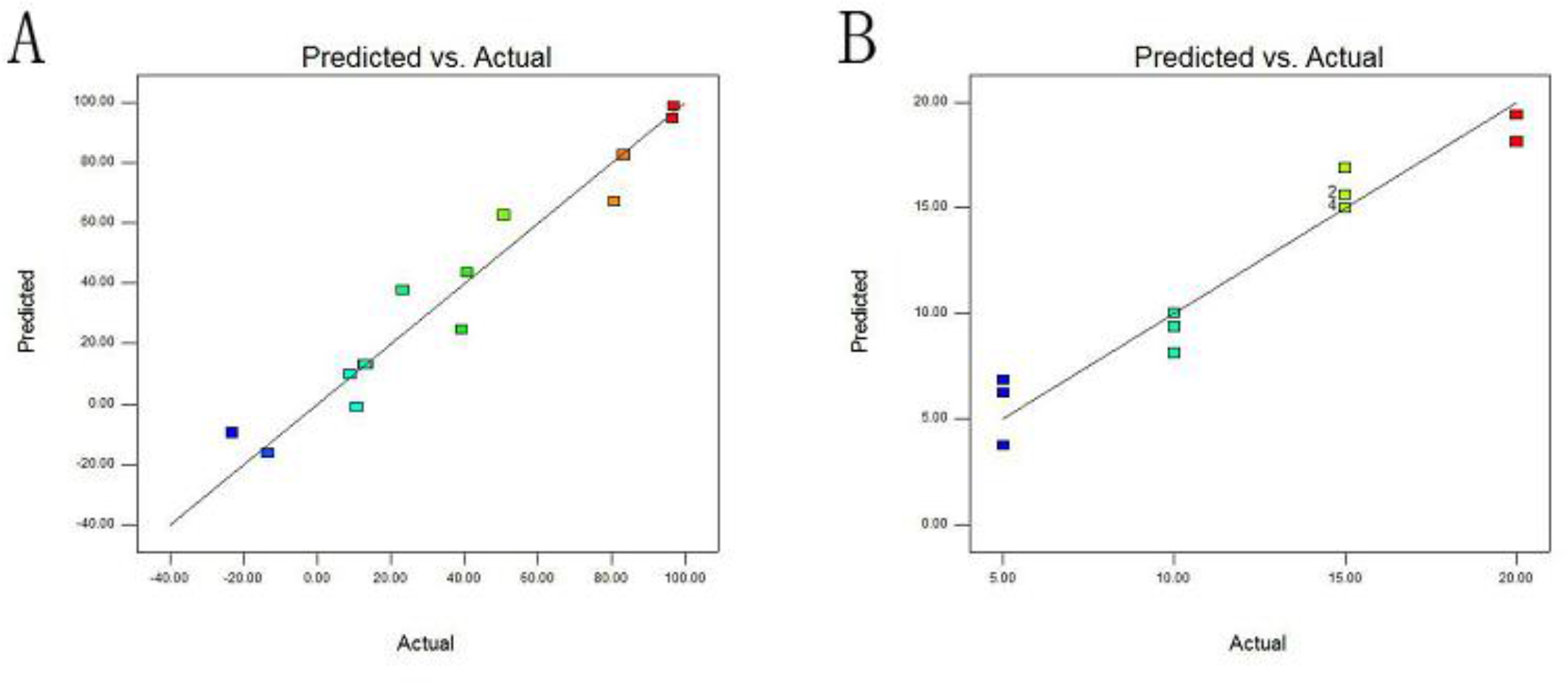
Plot of predicted compared with actual responses of (A) swelling degree and(B) membrane thickness.

**Tab 2.** ANOVA test on the quadratic regression models.

**Tab2 ANOVA test on the quadratic regression models**
| Response value | P-value | R<sup>2</sup> |
| --- | --- | --- |
| Y<sub>1</sub> | 0.0103* | 0.9476 |
| Y<sub>2</sub> | 0.0113* | 0.9454 |

3 Y<sub>1</sub>, swelling degree; Y<sub>2</sub>, membrane thickness.
p-value is the significant influence on the responses of the three factors.
\*Significant influence (p<0.05).

Three-dimensional response plot shown in Figures 2-3 showed the behavior of membrane thickness, swelling degree and integrity of membrane thickness, swelling degree and integrity rate, the main effect, interaction effect, and squared effect (nonlinear) of two independent variables at different levels. It was found that each of these three factors used in this study had its individual effect on the three responses. Fig. 2 showed that PLL concentration, volume ratio of PLL solution to beads and reaction time displayed a synergistic influence on S_w_, and the effect of reaction time was more significant than PLL concentration and volume ratio of PLL solution to beads. All the three factors exhibited the same effect just on membrane thickness as shown in Fig. 3.

**Fig. 2.**
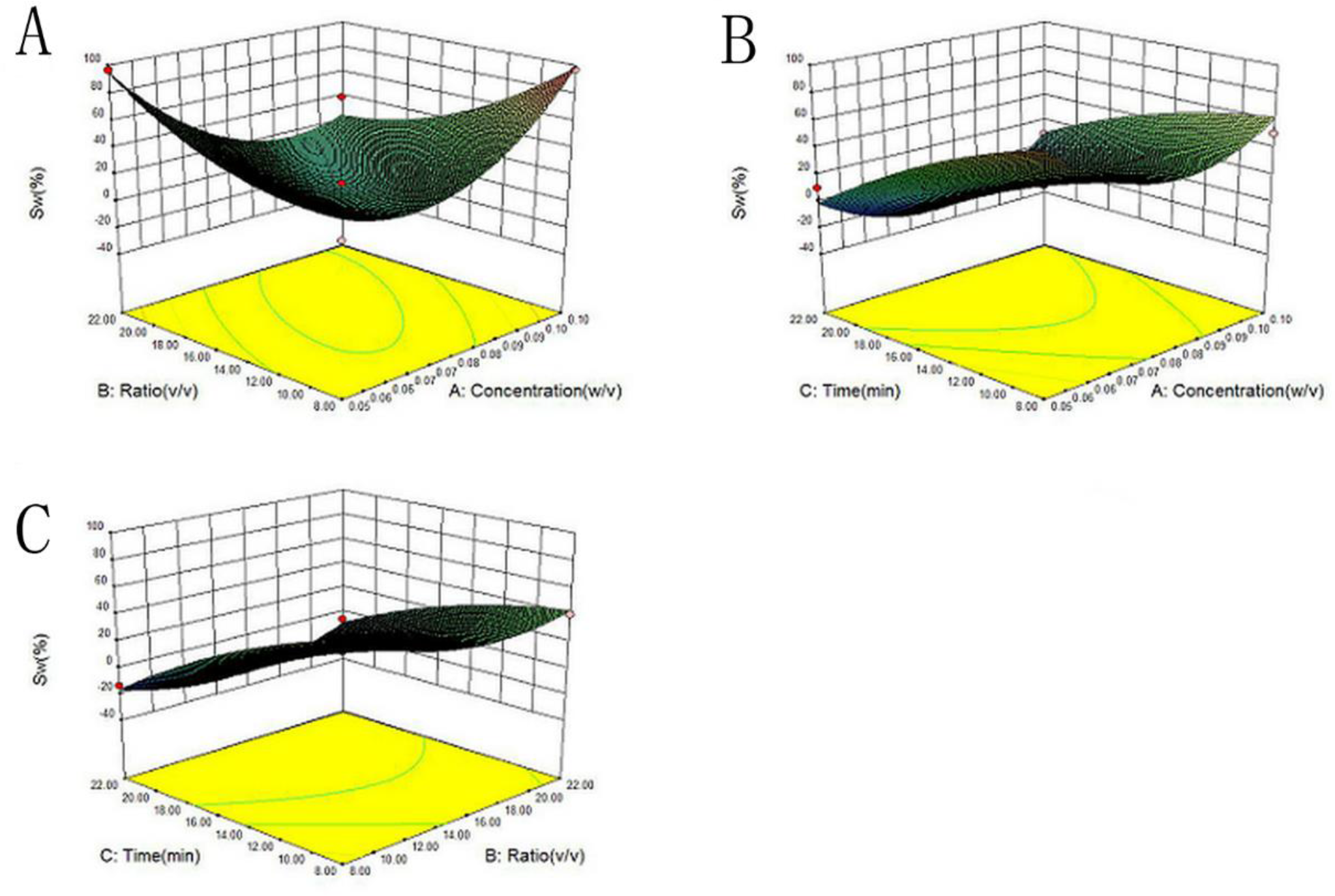
Response surface plots showing combined effect of (A) PLL concentration and volume ratio of PLL solution to beads, (B) reaction time and PLL concentration, and (C) reaction time and volume ratio of PLL solution to beads on S_w_.

**Fig. 3.**
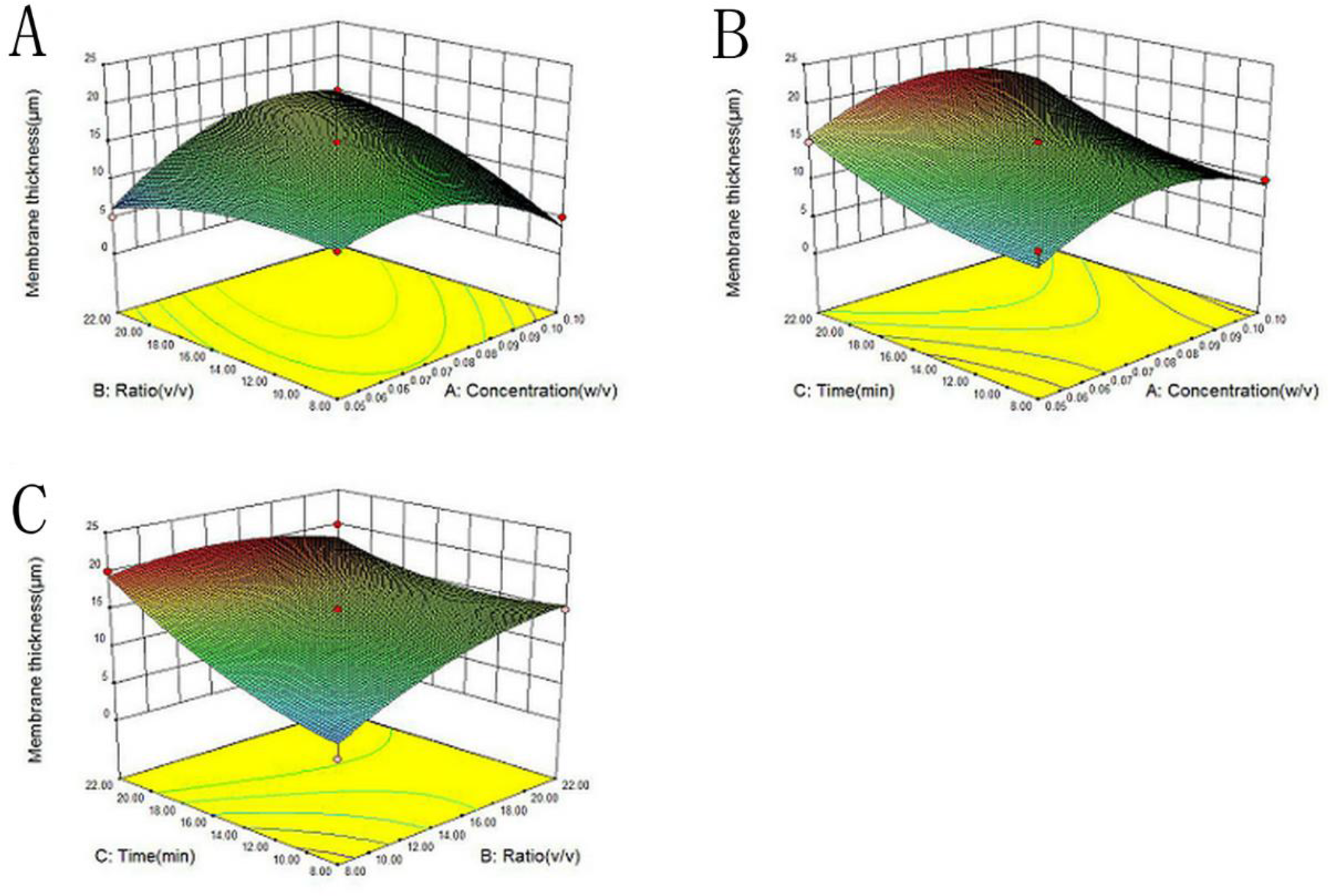
Response surface plots showing combined effect of (A) PLL concentration and volume ratio of PLL solution to beads, (B) reaction time and PLL concentration, and (C) reaction time and volume ratio of PLL solution to beads on membrane thickness.

### Optimum processing for maximum of membrane strength

Considering most suitable membrane thickness for cell culture (Ma Y et al. 2013), 15 μm was chosen as specified response and optimum processing for minimum of S_w_ was obtained by Box-Behnken model. Optimum conditions were X1=0.05, X2=15, X3=21.4. The Comparison of experiment value and predicted value was shown in Tab3.

**Tab 3.** Comparison of experiment value and predicted value.

| | $S_w$ (%) | |
| --- | --- | --- |
| Measured value | $3.54 \pm 1.02$ | |
| Predicted value | 3.89 |  |
| Agree (%) |  | 91.11 |

### Optimized microenapsules promote CHO cell viability and DSPA production

Microencapsulated CHO cells were made separately according to the optimum process referred above and literature (Huang et al., 2010). The membrane thickness in both groups were 15 μm, which was identified suitable for CHO cell growth. S_w_ of un-optimized microencapsules was 26.80±0.05%, while it was optimized to 3.54 ±1.02%. Optimized and un-optimized microencapsulated CHO cells were cultured in stirred tank bioreactor for 15 days. The morphology of microencapsulated CHO cells was observed under a light microscope (Fig. 4). It was found that broken microencapsule in un-optimized group on day 15, while none broken in optimized group. As shown in Fig. 5, both the proliferation and DSPA production of optimizated group were higher in the culture un-optimized. Membrane protect cells from destroyed, especially in long-term stir shear force. Unbroken microencapsule supply cells with stable environment for cell growth and activities. Thus, as expected, optimized microencapsules promoted the proliferation of microencapsulated CHO cells.

**Fig. 4.**
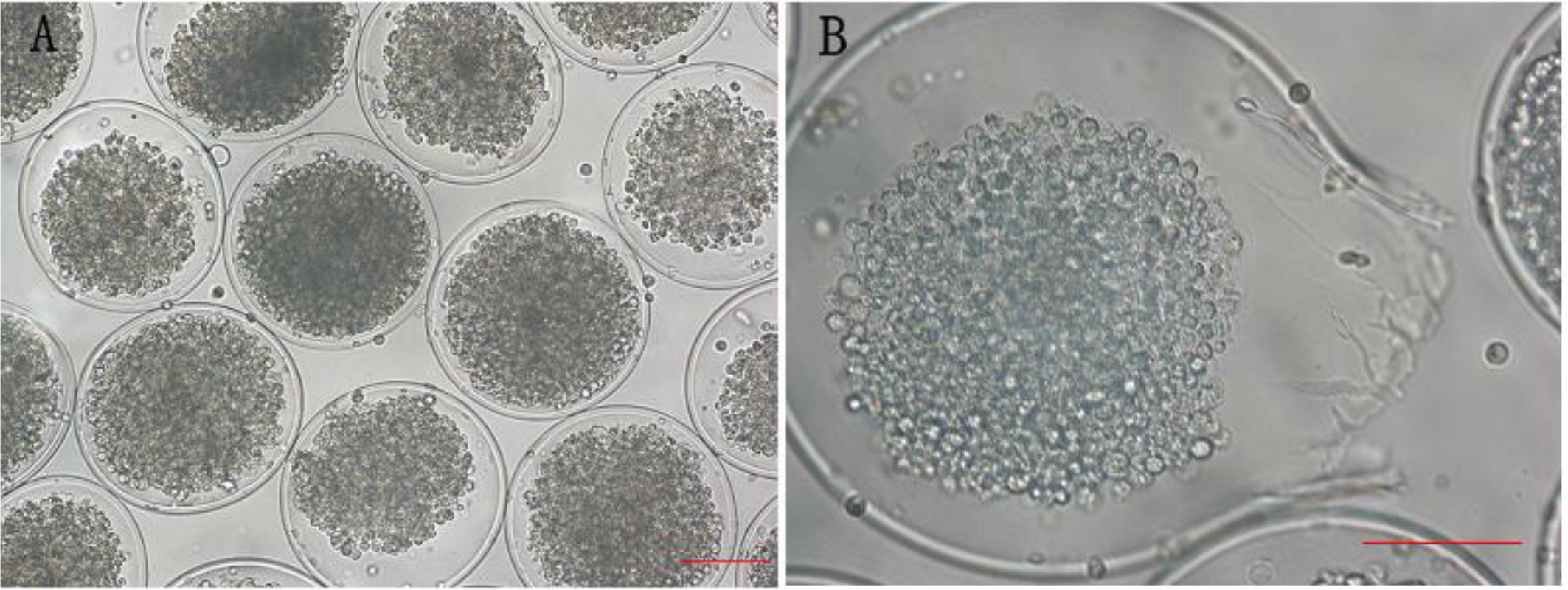
Growth profile of CHO cells cultured within microcapsules optimized and un-optimized. (A) Cells were cultured for 15 days within optimized microcapsules. (B) Cells were cultured for 15 days within un-optimized microencapsules, the membrane was broken. Scale bar = 100 μm.

**Fig. 5.**
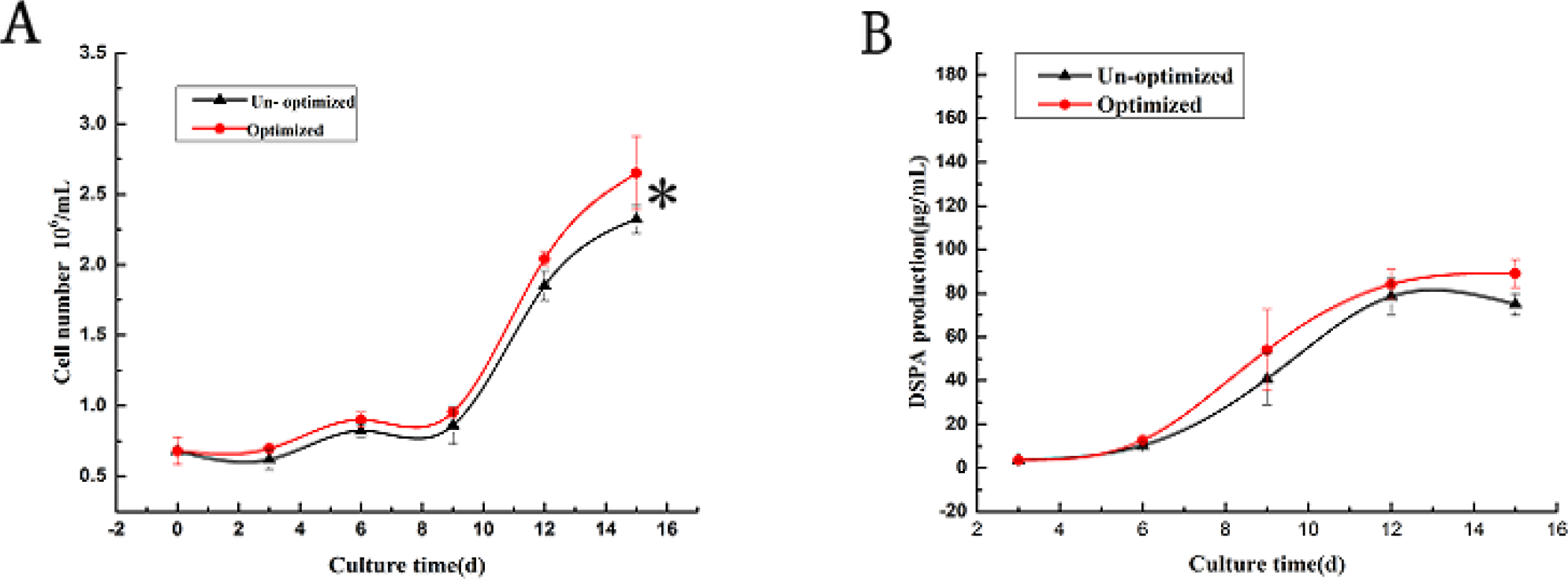
Comparison of CHO cells growth and protein production between optimized and un-optimized microencapsules. (A) Cell growth curve, “*” indicates statistically significant between the groups. (B) Protein production curve.

Cells were also examined by live/dead staining and observed under the confocal laser scanning microscope during the culture period, where the green fluorescence showed the calcein AM-labeled viable cells, and the red fluorescence showed the ED-1-labeled nonviable cells (Fig. 6). The strong green fluorescence and low-red fluorescence microcapsules indicated that both the microencapsulated cells remained high viability after preparation (Fig. 6 A1, B1). During subsequent culture, the two groups of microencapsulated cells obtained cell clusters, most of which were labeled by strong green fluorescence (Fig. 6 A2, B2). The results revealed that cells remained viable indicating that microcapsules with 15 um membrane thickness support cells proliferation in both groups.

**Fig. 6.**
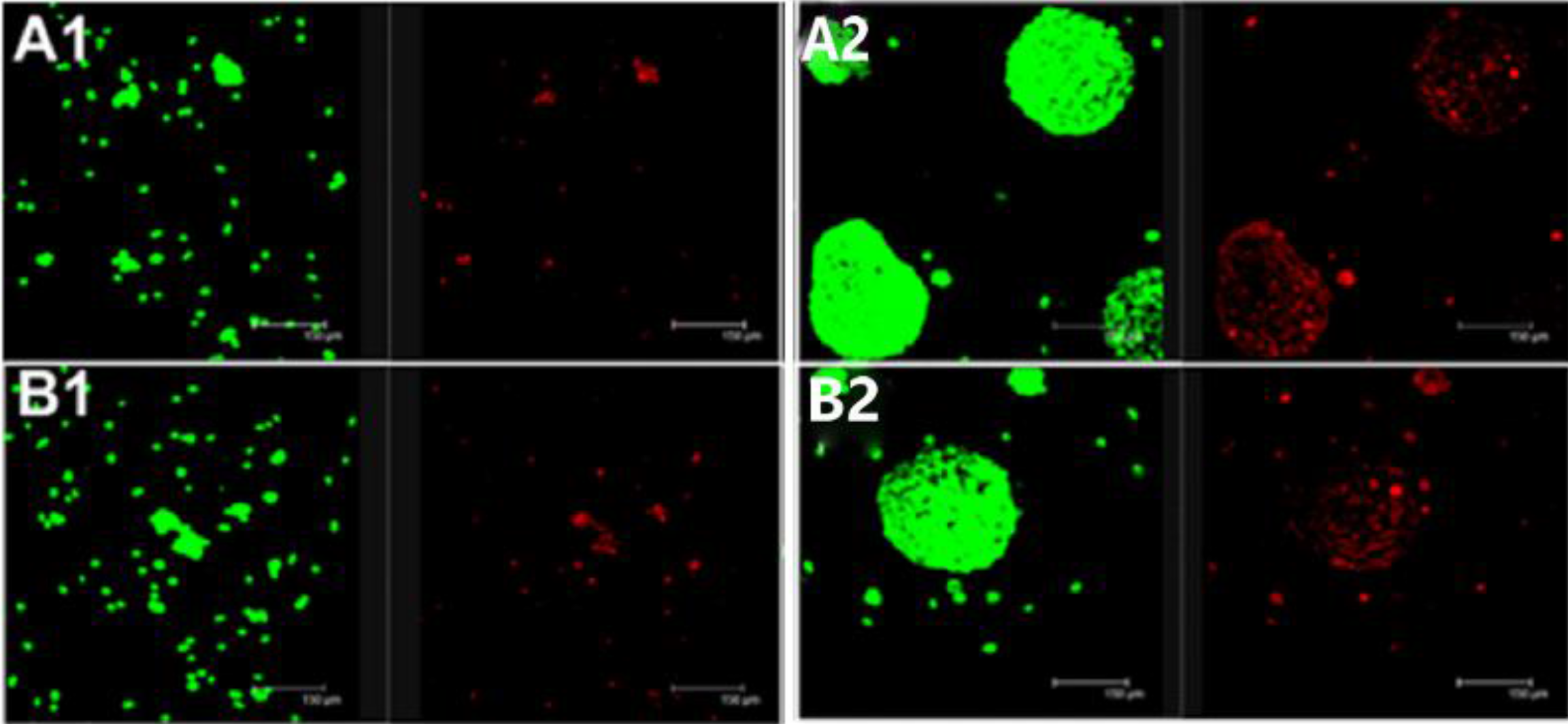
Live/dead staining of encapsulated CHO cells cultured for 3days and 21 days (green indicates live cells; red indicates dead cells). (A1, A2) Optimized microencapsules for 3 days and 21 days. (A1, A2) Un-Optimized microencapsules for 3 days and 21 days.

## Discussion

Results of ANOVA test on the quadratic regression models, presented in Tab. 3, indicated that responses were significant and adequate. Moreover, the results of model summary showed the R^2^ value was more than 0.9 for both the response, indicating a good correlation between the experimental and predicted responses and reliability of these models. Both of adequate precision ratios indicated an adequate signal. Reaction time showed higher effect on S_w_ and membrane thickness than others factors. It indicated that the reaction amount of PLL was mainly determined by reaction time.

The percentage prediction agreement for S_w_ indicated the robustness and high prognostic ability of the Box-Behnken models. On linear correlation of predicted and observed response variables, a correlation coefficient, R^2^ above 0.9, was got for membrane thickness and swelling degree. Thus, the higher magnitude of percentage prediction agreement (91.11%) as well as significant values of R^2^ in this study indicated the robustness and high prognostic ability of Box-Behnken models.

We could conclude that the optimized microencapsules with 15 μm membrane thickness and maximum strength were suitable for cell viability and long-time stirring. Although live/dead assay of CHO cells in un-optimized microencapsules appeared similar to optimized group, part of microencapsules were destroyed by stirring, result in cells leaking from microencapsules. CHO cells without protection from microencapsules would be damaged, or even die, that was why cell density in un-optimized microencapsules lower than optimized group in long-time stir.

In this experiment, a numerical optimization technique by using Box-Behnken model was used to develop a new formulation optimizing membrane strength with the desired membrane thickness. The verification result shows that Box-Behnken model is an efficient method to optimized process conditions of microencapsules for long-term cell culture.

## Materials and methods

### Preparation of microcapsules

Microcapsules were prepared as described previously (Huang et al., 2010). According to the literature, the concentration of α-poly-L-lysine (α-PLL), membrane formation time, and volume ratio of α-PLL solution to beads were chosen as three factors significantly affecting the membrane strength and thickness. The experiment was designed according to three factors and three levels Box-Behnken design by Minitab (Tab 4), X_1_-X_3_ was separately denoted as α-PLL (α-poly-L-lysine) concentration, the volume ratio of α-PLL solution to beads, and reaction time. 1.5% (w/v) sodium alginate solution(100 cp, the Chemical Reagent Corp, Qingdao, China), which was dissolved in 0.9% (w/v) NaCl solution, was extruded into 100 mM CaCl_2_ solution to form beads using a high voltage electrostatic generator (Shenzhen Second People’s Hospital, Guangdong, China). After been had hardened for 30 minutes, beads were washed with CaCl2 solution. The beads were then immersed in 0.05-0.1 % (w/v) α-PLL (Sigma, USA) for 8-22 minutes to initiate formation of a membrane around cells, the volume ratio of α-PLL solution to beads was 8-22, and then suspended in 55 mM sodium citrate solution for 10 minutes to liquefy the alginate-Ca gel core.

**Tab 4.** Design of Box-Behnken Method.

**Tab4 Design of Box-Behnken Method**
| Reaction conditions | X<sub>1</sub>(%) | X<sub>2</sub> | X<sub>3</sub>(min) |
| --- | --- | --- | --- |
| 1 | 0.075 | 8 | 8 |
| 2 | 0.05 | 15 | 8 |
| 3 | 0.1 | 15 | 8 |
| 4 | 0.075 | 22 | 8 |
| 5 | 0.05 | 8 | 15 |
| 6 | 0.1 | 8 | 15 |
| 7 | 0.075 | 15 | 15 |
| 8 | 0.075 | 15 | 15 |
| 9 | 0.075 | 15 | 15 |
| 10 | 0.05 | 22 | 15 |
| 11 | 0.1 | 22 | 15 |
| 12 | 0.075 | 8 | 22 |
| 13 | 0.05 | 15 | 22 |
| 14 | 0.1 | 15 | 22 |
| 15 | 0.075 | 22 | 22 |

2 X<sub>1</sub>, α-PLL concentration; X<sub>2</sub>, the volume ratio of α-PLL solution to beads; X<sub>3</sub>, reaction time.

### Measurement of microcapsules membrane thickness

The thickness of microcapsules membrane under different preparation conditions was determined under an optical microscopy (Olympus CK 40).

### Measurement of microcapsules S_w_ (swelling degree)

Examined the diameters of beads and microcapsules under an optical microscopy (Olympus CK 40). Membrane strength can be characterized by S_w_ and mechanical stability (Liu et al., 2004). Because mechanical stability is detected by ball milling which might destroy microcapsule to fragments, taking difficult to count microcapsules. Thus, measurement of S_w_ is more feasible and chosen to characterize membrane strength, the maximum of membrane strength is represented by the minimum of S_w_.

The formula of the S_w_ was shown as follows:

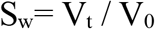

V_0_, the volume of a bead; V_t_, the volume of a microcapsule.

As beads and microcapsules were spherical, the formula could be expressed as follows (Liu X et al. 2004):

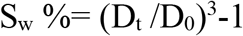

D_0_, the diameter of a bead; Dt, the diameter of a microcapsule.

### Validation of the model

15 μm was determined as the appropriate membrane for long-term cell culture (Ma et al., 2013), preparation of microcapsules was optimized for minimum of S_w_ according to the model with specific membrane thickness of 15 μm. Measured the S_w_ and compared with model prediction value.

### Long-term culture of the microencapsulated CHO cells

According to the verified formulation, microencapsulated CHO cells with maximum strength and 15 μm membrane thickness were prepared. As control, microencapsulated CHO cells with 15 μm thickness were prepared in experience according to literature. All microencapsules were cultured in stirred bioreactor with stirring rate of 90 rpm (Wang et al., 2015), which was a specific stirring rate identified to maximize the yield of recombinant protein. All of the microencapsulated cells were cultured in a growth medium consisting of CD OptiCHO medium supplemented with 4 mM glutamine. The volume ratio of microencapsulated cells to medium was 1:10, the medium volume was 50 mL, and the cells were grown at 37 °C under a 5% (v/v) CO_2_ atmosphere. The bioreactor was designed by the State Key Laboratory of Bioreactor Engineering, East China University of Science and Technology, Shanghai, People’s Republic of China.

## Detection and enumeration of microencapsulated cells

Detection and enumeration of microencapsulated cells based Cell Counting Kit-8 (CCK-8). The absorbance (A) of the supernatant was measured at 450 nm with measurement at 630 nm as a reference wavelength using a plate reader (Wellscan MK3; Thermo Labsystems, Helsinki, Finland). A standard curve was obtained for cell number against absorbance at 450 nm, using a range of dilutions of a known concentration of living cells as the standards. The formula for the relationship between living cell number (*Y*) and CCK-8 absorbance (*X*) was determined to be

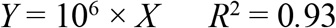

## Measurement of DSPA concentration

Measurement of DSPA concentration in the bioreactor media based on the fibrin plate assay to detect fibrinolysis activity (Whiting et al., 2014). Fibrin powder was added to an agar plate, providing a screening tool for plasminogen activator activity. We modified the assay as described previously (Whiting et al., 2014). Briefly, fibrinogen, prothrombin, and plasminogen (all from Sigma–Aldrich, St. Louis, MO, USA) were mixed with agar solution and left at room temperature for 30 Min for plate shaping. Standard concentrations of DSPA (10, 20, 40, 80, and 160 µg/mL) were added to the plates, which were then cultured at 37 °C for 5 H. DSPA degraded the fibrin and formed visible transparent lysis ring around each sample, and we measured the maximum and minimum diameters of every transparent loop. The DSPA concentration (denoted as C) had a logarithmic relationship to the mean diameter of transparent loop (denoted as D). The formula for the standard curve trend line was determined to be

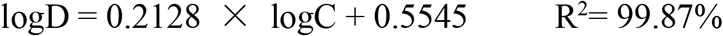

## Live/dead viability assay

Distribution of live and dead cells within microencapsulated cell clusters was detected by live/dead viability assay according to the protocol of manufacturer. Microencapsulated CHO cells were incubated with dual-color dyes (2 μM ED-1 and 1 μM calcein AM, sigma) at 37°C for 2 h. Then, the samples were scanned with CLSM (Leica SP2, Germany).

## Statistical analysis

All data were expressed as Mean ± SEM and Box–Behnken design was performed and statistically analyzed using Design Expert software V8.0.6.1 (Stat-Ease, Minneapolis, MN, USA). Statistical analysis at each sample point was calculated using ANOVA. A p-value < 0.05 was considered to be statistically significant (*).

